# ISG15 orchestrates dynamic crosstalk between mitochondrial fat oxidation and type 1 interferon in myeloid cells

**DOI:** 10.64898/2026.01.19.700051

**Authors:** Anand Kumar Gupta, Jing Wu, Pijush Das, Rahul Sharma, Sonali Das, Kim Han, Rachael J. Klein, Rebecca D. Huffstutler, Christian Combs, Warren J. Leonard, Claudia Kemper, Erin E. West, Michael N. Sack

## Abstract

In contrast to Krebs cycle intermediates, the role of mitochondrial fatty acid oxidation (FAO) in immunometabolism remains incompletely characterized. Studying primary bone marrow-derived macrophages (BMDMs), we show that IFN-β and STING activation augments FAO and associated enzymes carnitine palmitoyltransferase 1a (CPT1a) and acetyl-CoA acetyltransferase 1 (ACAT1). Depleting BMDM *Cpt1a* reduces FAO and dampens type 1 interferon (IFN) signaling due to decreased histone H3-K9/K14 acetylation, supporting the prior finding of an epigenetic role of FAO in sustaining type 1 IFN. Interestingly, FAO induction by IFN was dynamic as a heightened IFN response suppressed FAO, suggesting a concurrent negative-feedback mechanism. This FAO blunting correlated with increased expression of interferon-stimulated gene 15 (ISG15), a ubiquitin-like modifier known to modulate metabolic proteins through ISGylation. This immune-metabolic signature was similarly operational in mice infected with lymphocytic choriomenigitis virus (LCMV) with temporal discordance between ISG15 levels and FAO. The role of ISG15 in this negative feedback was shown with increased FAO and type 1 IFN response in *Isg15* knockout BMDMs. Parallely, endogenous co-immunoprecipitation showed interactions between ISG15 and CPT1a/ACAT1. This ISG15-FAO regulatory interaction was also evident in systemic lupus erythematosus (SLE)-associated interferonopathy and in primary monocytes from SLE individuals that exhibited increased ISG15 levels and reduced FAO rates. Collectively, these findings support a model in which type 1 IFN initially enhances FAO to amplify interferon production, but excessive IFN-signaling induces ISG15-mediated inhibition of FAO, a putative feedback loop that restrains inflammation and preserves immune homeostasis. Together these data identify a novel biphasic FAO-dependent immunometabolic regulatory program.

**Graphical Summary:** 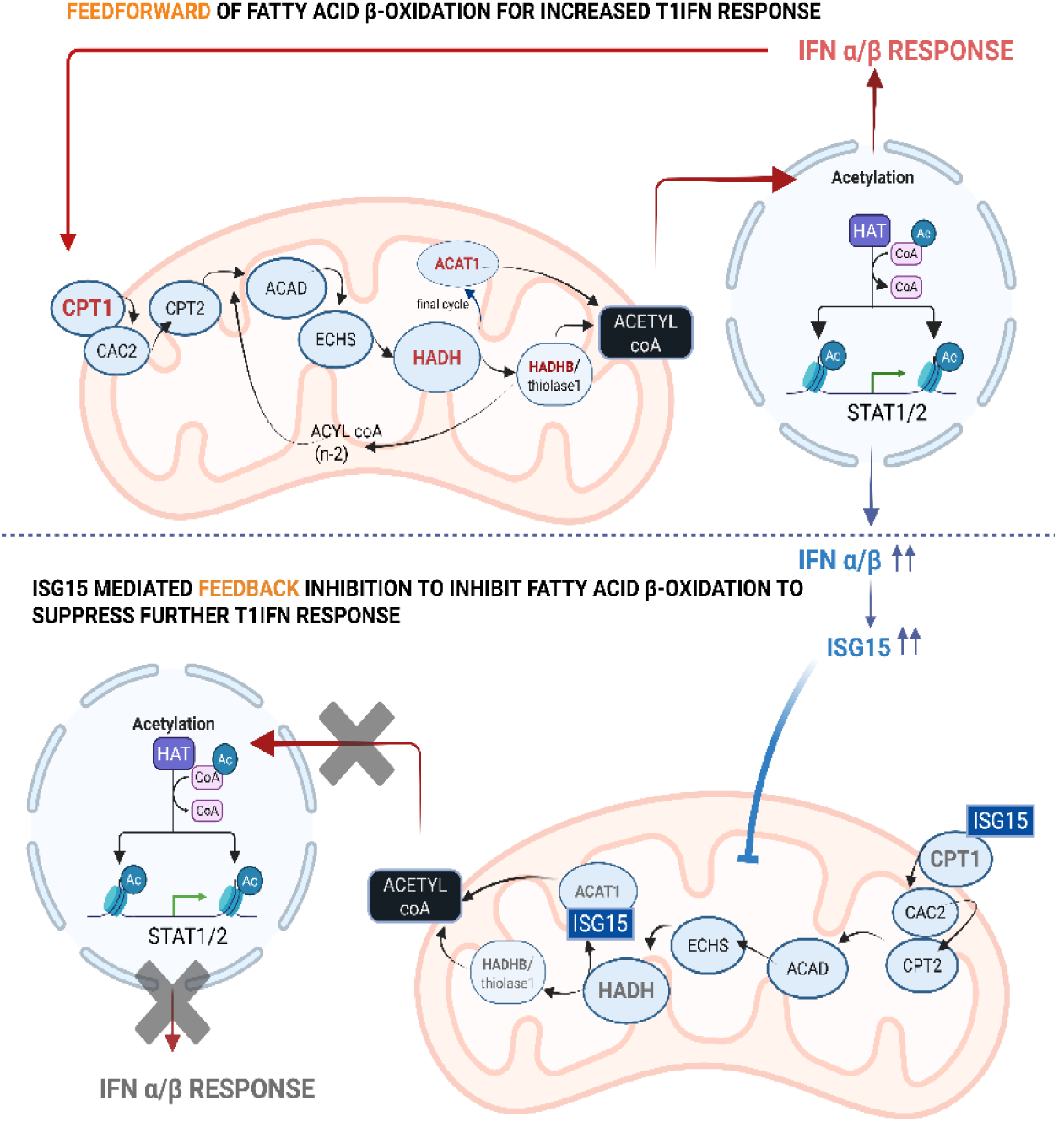

**Highlights:**

- Type 1 IFN rewires myeloid cell metabolism towards FAO.
- FAO epigenetically fuels Type 1 1IFN response.
- Effect of IFN on FAO is dynamic: inducing FAO at low concentrations while suppressing it during a heightened IFN response.
- Enhanced Type 1 IFN response produces ISG15 which ISGylates key enzymes of FAO altering their stability
- ISG15 negatively regulates FAO to control Type 1 IFN homeostasis in a negative feedback loop.

---

Cellular metabolism is critical in dictating the fate and functions of immune cells. This concept was initially investigated in the context of increasing metabolic demand by comparing quiescent cells to effects of immune cell differentiation, proliferation and activation.(1) Subsequent studies expanded our understanding of immunometabolism where individual metabolites were found to function as signaling molecules, serving as either G-protein coupled receptor (GPRC) signaling ligands or as substrates in post-translational control of regulatory proteins.(2–6) More recently, metabolites have been shown to confer epigenetic regulatory effects via their roles as substrates for acetylation, lactylation and/or methylation of DNA and histones.(6, 7)

To date, the most comprehensive studies on immunometabolism have been predominantly performed in the context of glucose, glutamine and ketone metabolism, and by investigating the effects of tricarboxylic acid cycle (TCA) intermediates.(1, 8–10) Our understanding of fatty acid metabolism in this regard is less well characterized, although it has widely implicated with immune cell plasticity and polarization. Alternatively activated, in contrast to classically activated macrophages have been reported to be dependent on mitochondrial fat oxidation for fueling their immune regulatory responses.(11) Fat metabolism was also important in the adaptive immune response, with mitochondrial fatty acid β-oxidation (FAO) being required for CD8^+^ T memory cell function and for CD4^+^ T regulatory cell proliferation and differentiation.(1) Additionally, short chain fatty acids serve roles as GPCR ligands, and in histone deacetylase inhibition in both innate and adaptive immune inflammatory pathways.(5, 12, 13). The crosstalk between fat metabolism and immune response is probably multidimenstional, illustrated in part, where NK derived IFN-γ initiates adipose tissue lipolysis to release free fatty acids which promote B cell activation in response to viral infection in mice.(14) The role of mitochondrial FAO in myeloid cell immunoregulation was initially described by studies showing that interferon-β (IFN-β), via peroxisomal proliferator-activated recepeptor-α (PPARα) activation, enhances FAO during plasmacytoid dendritic cell activation.(15) Similarly, another study showed an interplay between the metabolism of polyunsaturated fatty acids and the STING (stimulator of interferon genes)-dependent inflammatory responses.(16) Previous evidence from our laboratory shows that acetyl-CoA acetyltransferase 1 (ACAT-1), the penultimate enzyme of the FAO, which via acetyl-CoA, mediates the extent of histone acetylation on type 1 IFN genes, thus conferring an epigenetic regulatory role in monocytic type 1 IFN signaling.(7) Furthermore, other studies suggest a role of interferon-stimulated gene 15 (ISG15) in controlling myeloid cell lipid metabolism during viral infections,(17) although its role in FAO has not been characterized. Here, we have investigated the regulatory interactions between myeloid cell type 1 IFN and FAO, the effect of ISG15 on FAO, and whether these regulatory programs play an important role in myeloid cell immunometabolism and immunoregulation.

In this study we demonstrate that type 1 IFN rewires myeloid cell fat metabolism in a dynamic fashion employing different regulatory control nodes. The initial type 1 IFN response induces mitochondrial FAO so as to epigenetically promote a heightened type 1 IFN response in a feedforward manner. However, at higher concentrations or prolonged activation with type 1 IFN or STING inhibits mitochondrial FAO. We further show that ISG15, a ubiquitin-like modifier, functions as the negative molecular switch by ISGylating FAO enzymes, thereby controlling FAO to restrain further type 1 IFN activation. Together, this early epigenetic control and the delayed role of ISG15 appear to function in concert as novel immunometabolic modulators to integrate type I interferon signaling and mitochondrial FAO.

## Results

### Type I interferon signaling upregulates mitochondrial FAO in myeloid cells

To study the interplay between FAO and type 1 IFN we initially employed a STING agonist, DMXAA ((ASA 404) (18)), which showed a time-dependent induction of the type 1 IFN response, with maximal induction at 2 hours (Supplementary Figure 1A). The effect of this agonist was then used to determine the effect on genes encoding FAO enzymes and on corresponding protein levels in primary murine bone marrow derived macrophages (BMDMs). Figure 1A is a schematic highlighting key enzymes linked to uptake of long chain fatty acids and involved in mitochondrial FAO. Compared to vehicle control, BMDMs supplemented with DMXAA for 2 or 4 hours showed induction of transcript levels of FAO encoding genes (Figure 1B). More specifically, the transcripts encoding the trifunctional protein hydroxyacyl-CoA dehydrogenase alpha subunit (*Hadha*) was significantly induced at 2 hours, and the transcripts encoding carnitine palmitoyltransferase 1A (*Cpt1a*) and *Acat1* were significantly elevated after 4 hours of DXMAA exposure (Figure 1B). CPT1 is rate limiting in FAO as it is responsible for the transport of long chain fatty acids, the major oxidative phosphorylation intermediates, into the mitochondrial matrix. To evaluate whether BMDM oxidative phosphorylation and FAO were modified by DMXAA, the basal and maximal respiration were assayed in the presence or absence of the CPT1 inhibitor etomoxir. Consistent with prior data using cGAMP as a type 1 IFN trigger,(19) DMXAA and IFN-β increased basal and maximal mitochondrial oxygen consumption rates (OCR) (Figure 1C-E, Supplementary Figure 1B-D). This effect on OCR was confirmed using IFN-β and DMXAA as the stimuli in BMDMs where palmitate was used as the major mitochondrial substrate in low glucose media. Both stimuli increased OCR relative to vehicle treated control cells and etomoxir blunted both basal and maximal respiration following IFN-β stimulation (Figure 1F-K). We additional assessed FAO, by quantifying FAOBlue fluorescence. Mitochondrial oxidation of this coumarin derivative bound to a 9-carbon chain fatty acid, results in its catabolism to propionic acid. This releases coumarin into the cytoplasm resulting in a strong blue fluorescence excitation signal that reflects the rate of FAO (Figure 1L). Here, and consistent with the OCR data, FAO was significantly induced by both DMXAA and INF-β (Figure 1M-N). Collectively, these data show that type I interferon triggers rewire or drive myeloid cells metabolism towards mitochondrial FAO.

**Figure 1:**
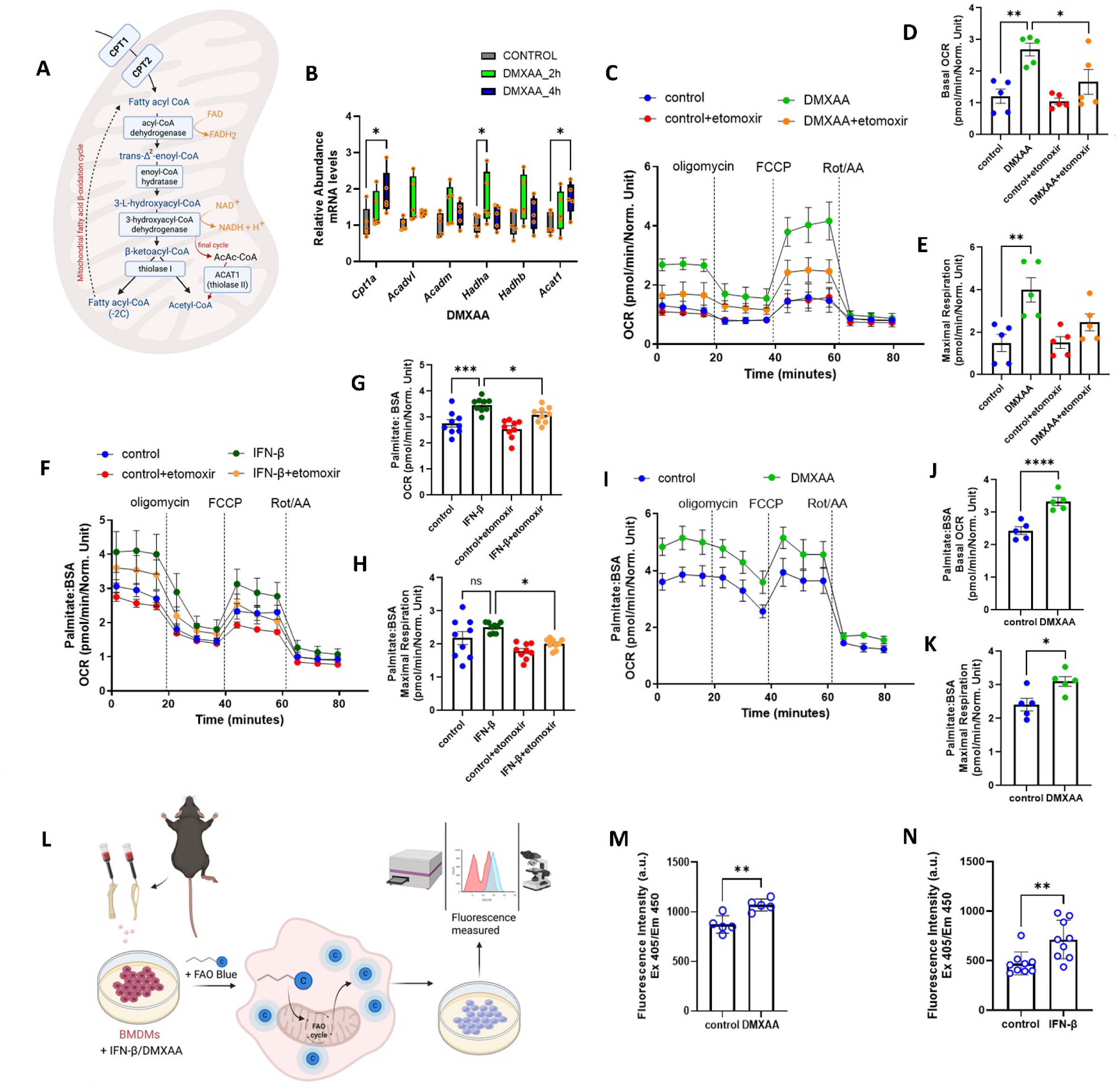
Type I interferon response reprograms myeloid cell metabolism towards mitochondrial fat oxidation. **(A)** Schematic representation of mitochondrial fatty acid β-oxidation pathway. **(B)** Quantitative RT-PCR analysis of key enzymes of FAO in BMDMs stimulated with or without 10 μg/mL DMXAA for 2h and 4h respectively (n=3 per group). Data normalized to 18*S* rRNA and represented as means ± SEM. Two-way ANOVA followed by Tukey’s multiple comparisons test. **(C)** Seahorse analysis of oxygen consumption rate (OCR) of BMDMs stimulated with 10 μg/mL DMXAA for 4 h with or without etomoxir pre-treatment treatment (n=5 per group). Graphical representation of basal OCR **(D)** and maximal respiration **(E)** of Seahorse analysis in C. **(F-K)** Seahorse analysis of OCR along with respective graphical representations of basal OCR and maximal respiration of IFN-β **(F-H)** (n=9 per group) and DMXAA **(I-K)** (n=5 per group) treated BMDMs in presence of 200µM palmitate:BSA as the substrate. Measurement of OCR of BMDMs **(C-K)** were done after sequential treatment of oligomycin, FCCP and Rotenone/Antimycin A. Results measured were normalized to the cell counts in each corresponding wells. Two-way ANOVA followed by Tukey’s multiple comparisons test. **(L)** Schematic representation of using FaO Blue as an alternative approach to measure mitochondrial fat oxidation. **(M-N)** Histograms of FaO Blue intensity measured at Excitation 405/ Emission 450 in BMDMs treated with DMXAA **(M)** (n=5 per group) and IFN-β **(N)** (n=9 per group) for 4h. Fluorescent intensity were normalized to the cell counts and represented as means ± SEM using an unpaired t Test, with dots representing n per group. Asterisks represent - \**p* < 0.05; \*\**p* < 0.01; \*\*\**p* < 0.001. n.s. - not significant.

### Disruption of mitochondrial fatty acid uptake perturbs type 1 IFN induction of oxidative phosphorylation and blunts inflammatory signaling

To validate the interplay between FAO and type 1 IFN, the carnitine-dependent transport of fatty acids across the mitochondrial membrane was genetically disrupted in primary BMDMs using CRISPR-Cas 9 mediated targeting of *Cpt1a* as shown (Figure 2A) resulting in an ≈ 90%. reduction in primary BMDM Cpt1 steady-state levels (Supplemental Figure 2A-B). We then evaluated the rate of palmitate driven OCR in response to *Cpt1a* knockout (KO). In the presence of DMXAA for 4 hours, maximal but not basal respiration was significantly attenuated in *Cpt1a* KO BMDMs (Figure 2B-D). In parallel, the secretion of IFN-β was blunted in *Cpt1a* KO BMDMs compared to controls (Figure 2E). Interestingly, TLR4 activation by lipopolysaccharide exposure similarly reduced IFN-β release from *Cpt1a* KO BMDM’s at 4 hours of stimulation (Supplemental Figure 2C). In parallel, DMXAA-induced steady-state and phosphorylation levels of STAT1, STAT2 and IRF3, three key mediators of type 1 IFN induction, were all reduced in the absence of Cpt1a (Figure 2F and Supplemental Figure 2D). Interestingly, ISG15 levels were similarly blunted by *Cpt1a* KO mediated reduction in FAO capacity (Figure 2F and Supplemental Figure 2D). In line with numerous levels of regulation we had previously shown that ACAT1 knockout blunted type 1 IFN signaling via epigenetic regulation (7). Correspondinlgy, acetylation of His3 K9/K14 was blunted in *Cpta1* KO BMDMs in response to DMXAA stimulation (Figure 2G and Supplemental Figure 2E), which reinforces an acute epigenetic regulatory role of FAO in modulating type 1 IFN biology (Figure 2H).

**Figure 2:**
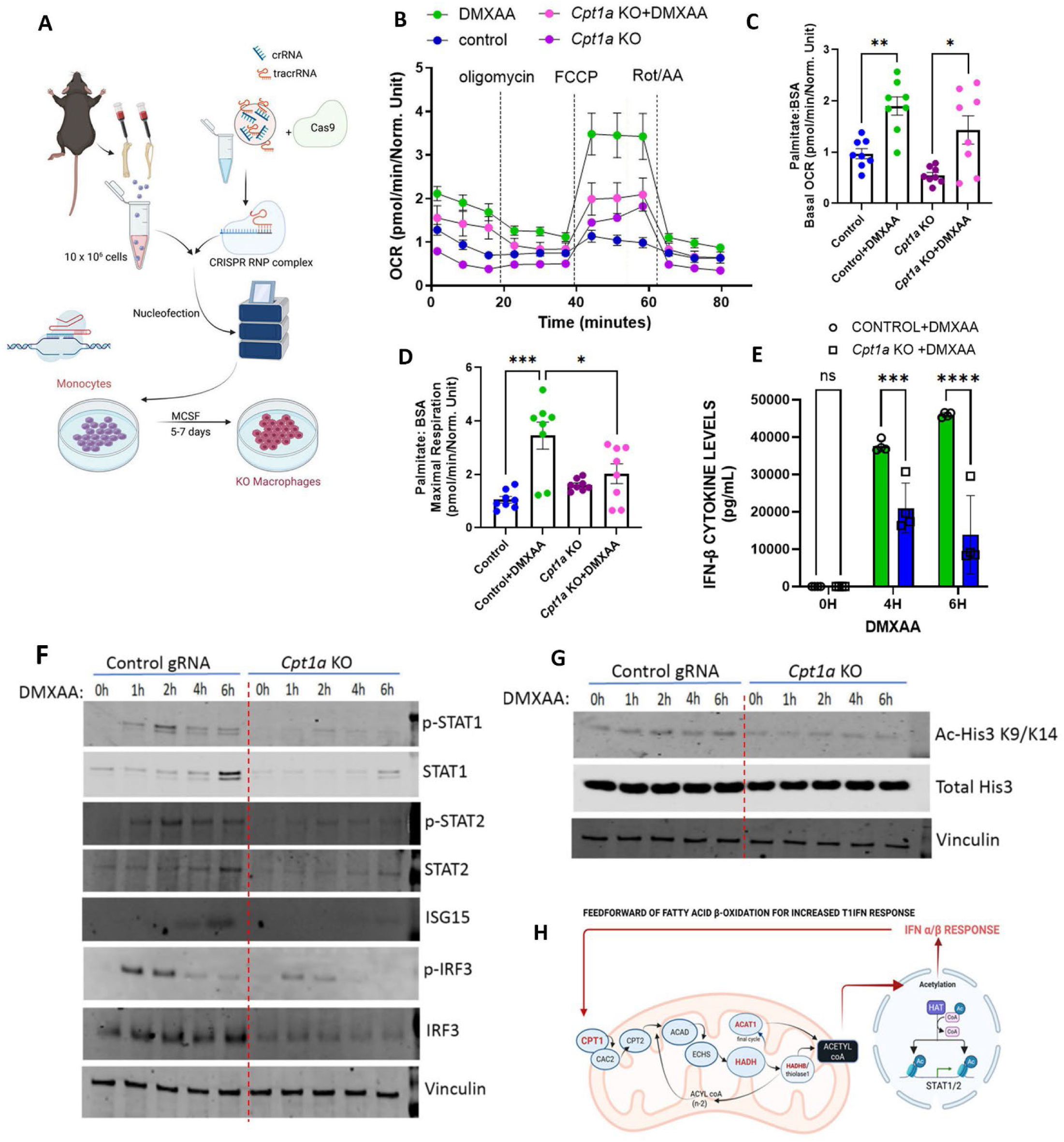
Blocking mitochondrial FAO blunts the type 1 IFN response. **(A**) Strategy for CRISPR Cas9 mediated genetic depletion of *Cpt1a* in primary BMDMs. **(B)** Seahorse analysis of OCR in *Cpt1a* KO and control BMDMs on DMXAA stimulation on sequential treatment with oligomycin, FCCP and Rotenone/Antimycin A (n=8 per group). **(C-D)** Histograms of basal OCR **(C)** and maximal respiration **(D)** of seahorse analysis in **B**. **(E)** Histograms showing IFN-β cytokine levels in *Cpt1a* KO and control BMDMs (n=4 per group) on treatment with DMXAA for indicated time points (0-6h). Results measured were normalized to the cell counts in each corresponding wells using two-way ANOVA followed by Tukey’s multiple comparisons test. **(F)** Representative immunoblots of pSTAT1, STAT1, p-STAT2, STAT2, ISG15, p-IRF3, IRF3 and vinculin in DMXAA treated control and *Cpt1a* KO BMDMs upon treatment with DMXAA for indicated time points (0-6h) (n=3 per group). **(G)** Representative blots of Ac-His3 K9/K14, total His3 and vinculin in control and *Cpt1a* KO BMDMs upon treatment with DMXAA for indicated time points (0-6h) (n=3 per group). **(H)** Schematic of proposed epigenetic control whereby type 1 IFN in a feed forward manner increases further interferon production in response by increaing myeloid cell mitochondrial FAO to generate Acetyl-CoA and histone acetylation. Asterisks represent - \**p*< 0.05; \*\**p* < 0.01; \*\*\**p*< 0.001. n.s. - not significant.

### The link between FAO and type 1 IFN is evident in LCMV infection

To explore the relationship between the regulation of FAO and type 1 IFN *in vivo*, we employed a murine lymphocytic choriomeningitis virus (LCMV) infection as a canonical antiviral response model of acute infection induced monocyte/macrophage-derived type 1 IFN.(20) Wildtype C57BL/6 mice were infected with LCMV via tail vein injection (Figure 3A) and sacrificed at days 1, 3, 5 and 8 (n = 5 for each timepoint). Splenic CD11b^+^ cells were isolated by positive selection. Transcript levels were assessed by RT-qPCR, and mRNAs encoding *Stat1*, *Stat2* and *Ifnb* peaked at day 3, whereas the transcript levels of the gene encoding *Isg15*, was delayed and peaked at day 5 (Figure 3B-E). In parallel, levels of genes encoding the FAO pathway proteins HADHA and the β-subunit HADHB peaked at day 1 (Supplementary Fig. 3A-B), and *Cpt1a* and *Acat1 mRNAs* peaked at day 3 (Figure 3F-G). The functional effects on FAO by LCMV infection was determined in CD11b+ myeloid cells from the splenocytes using FAOBlue fluorescence by confocal microcopy and flow cytometery.(21) Both techniques showed increased FAO at day 1 and day 3 post infection with gradually reducing FAO rates towards basal levels by day 8 (Figures 3H-K). Collectively, *in vivo* LCMV infection resulted in similar patterns of induction in type 1IFN and FAO as evident in the BMDMs. Furthermore, induction of *Isg15* mRNA was delayed and peaked as FAO enzyme transcript levels were diminishing. These data suggest that the FAO-type 1 IFN axis is dynamic and the question arises as to how this interaction is regulated?

**Figure 3:**
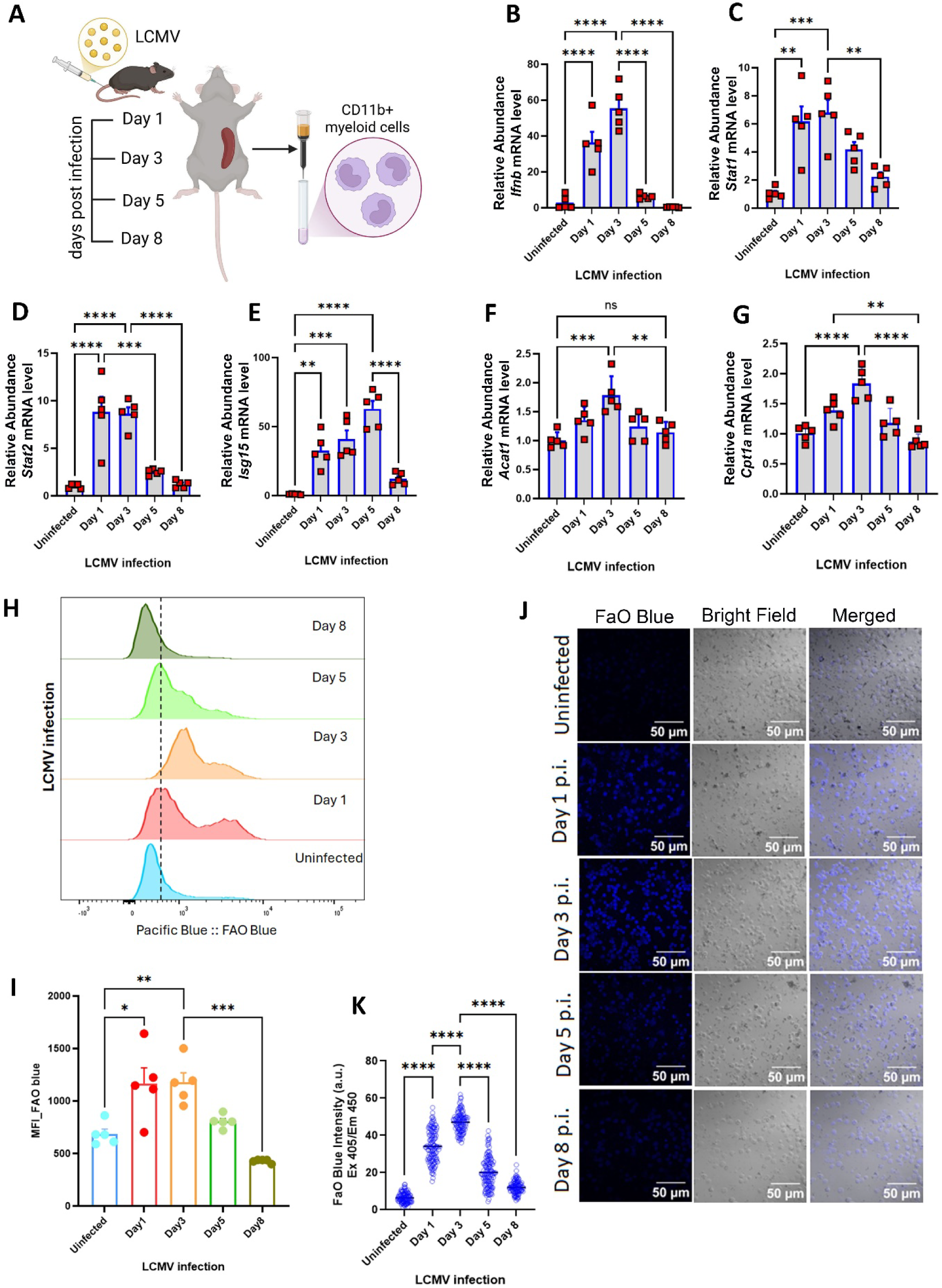
Crosstalk of mitochondrial FAO and type 1 IFN in myeloid cells during *in vivo* LCMV infection. **(A**) Schematic of the murine infection study design. LCMV infected C57bl/6 mice were sacrificed at indicated time points (0,1,3,5 and 8 days) post infection and splenocytes were isolated (n=5 mice per group/day). CD11b+ myeloid cells were separated from the splenocytes and used for this study. **(B-G)** Histograms showing quantitative RT-PCR analysis of type 1 IFN response genes *Ifnb* **(B)**, *Stat1* **(C)**, *Stat2* **(D)**, *Isg15* **(E)** and mitochondrial FAO enzymes *Acat1* **(F)** and *Cpt1a* **(G)** in CD11b+ myeloid cells isolated from the splenocytes of infected mice. Data were normalized to 18*S* rRNA and represented as means ± SEM. Splenocytes from infected mice were subjected to treatment with 5µM FaO Blue followed by staining with PE conjugated CD11b+ antibody. **(H)** Representative data of flow cytometry based measurement of fluorescence intensity of FaO Blue in CD11b+ splenocytes from infected mice at indicated time points **(J)** Representation of flow cytometric geometric mean fluorescence intensity of FaO Blue in infected mice at different timepoints. **(K)** Representative microscopic images of CD11b+ myeloid cells, isolated from the splenocytes of infected mice at different timepoints, incubated with FaO Blue. **(I)** Histograms of fluorescence intensity of FaO Blue from microscopic images in **I**. Data represented as means ± SEM. Two-way ANOVA followed by Tukey’s multiple comparisons test. Asterisks represent \**p* < 0.05; \*\**p* < 0.01; \*\*\**p* < 0.001. n.s. - not significant.

### The interplay between type 1 IFN and FAO is dose- and time-dependent

To uncover the temporal dynamics of FAO with type 1 IFN post infection, we compared the oxygen consumption rate in BMDMs at later time points of DMXAA exposure compared of the initial 4 hours of stimulation described previously. In contrast to the acute stimulation, the rate of FAO was reduced following 24 hours of STING activation compared to the shorter duration of stimulation (Figure 4A-C). We hypothesised that the type 1 IFN response may have a dynamic effect on FAO, i.e., inducing FAO during a modest stimulatory response in contrast to FAO inhibition during heightened or prolonged type 1 IFN stimulation. To test this concept, BMDMs were then subjected to different concentrations of IFN-β for 4 h. IFN-β at 50 ng/mL robustly induced FAO, whereas doses of 100 and 200 ng/mL reduced palmitate-driven FAO compared to the 50ng/mL IFN-β exposure (Figure 4D-F). Interestingly, this same dose-dependent biphasic pattern was evident at the transcript levels encoding *Cpt1a* and *Acat1* (Figure 4G-H). In contrast, the transcript levels encoding *Isg15* increased incrementally with higher doses of IFN-β exposure (Figure 4I). This discordance resulted in a negative correlation between the transcript amounts of *Isg15* and *Cpt1a* and *Acat1* respectively (Figure 4J-K). Moreover, a negative correlation between *Isg15* levels and palmitate driven oxygen consumption rate was also observed (Supplementary Figure 4A-B). The viability of BMDMs were also assessed in presence of DMXAA for longer durations (up to 24 hours) and at higher concentrations of IFN-β. BMDM viability was not perturbed by STING activation for this duration or higher dose (upto 200 ng/mL) of IFN-β for 4 h (Supplementary Figure 4C-D).

**Figure 4:**
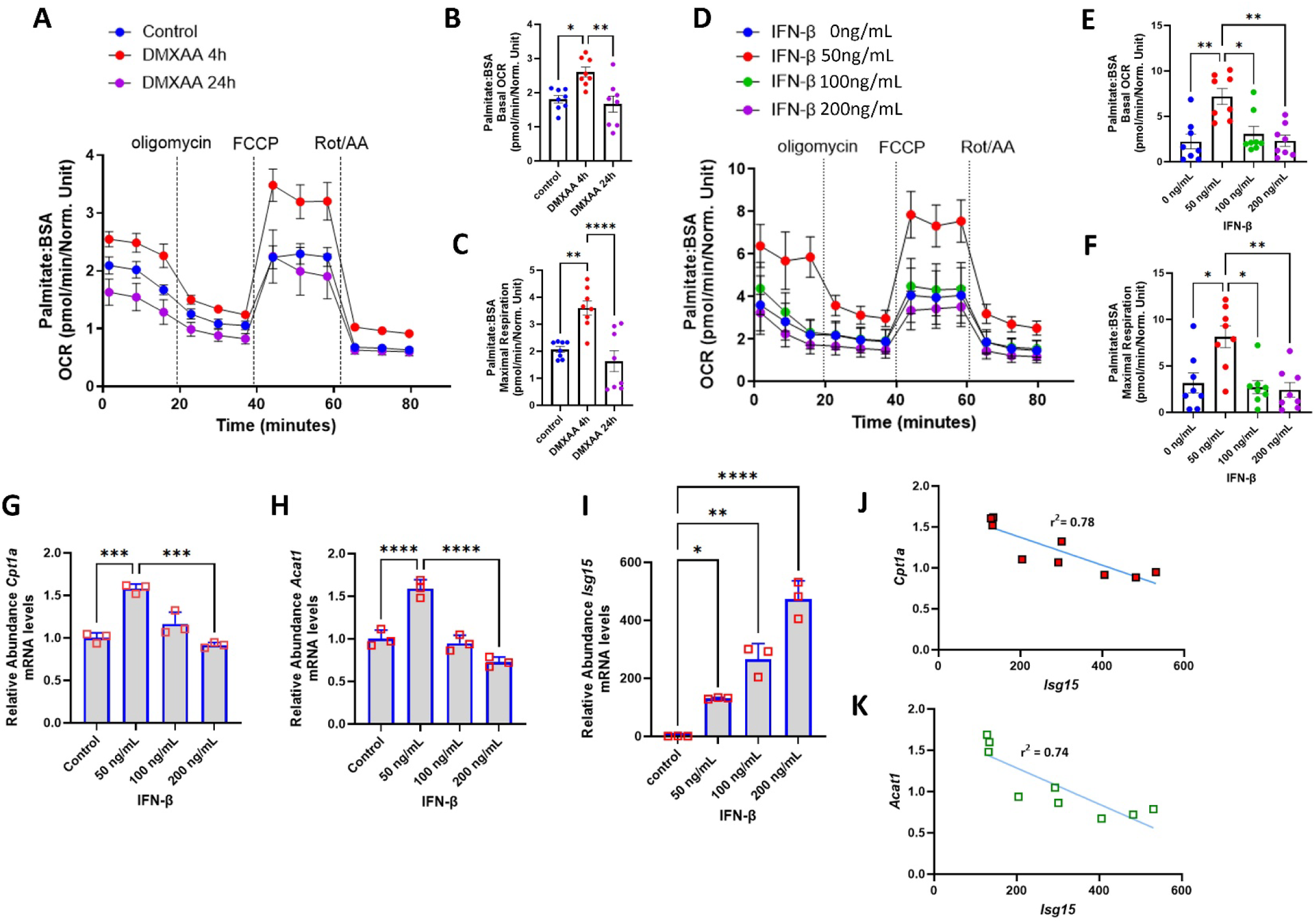
Temporal and dose dependent dynamics of myeloid cells FAO in response to type 1 IFN signaling. **(A**) Seahorse analysis of BMDMs upon stimulation by DMXAA for different time periods (0-24h) in presence of palmitate:BSA after sequential treatment with oligomycin, FCCP and Rotenone/Antimycin A (n=8 per group). Histogram representing basal OCR **(B)** and maximal respiration **(C)** from all replicate Seahorse experiments described in panel **A. (D)** Seahorse analysis of BMDMs upon stimulation by increasing doses of IFN-β (0 ng/mL to 200ng/mL) (n=8 per group). **(E-F)** Histograms of basal OCR **(E)** and maximal respiration **(F)** of BMDMs in response to different IFN-β doses. Results measured were normalized to the cell counts in each corresponding wells. **(G-I)** Histograms of quantitative RT-PCR analysis of *Cpt1a* **(G)**, *Acat1***(H**) and *Isg15* **(I)** expression in myeloid cells stimulated with indicated concentrations of IFN-β (n=3 per group). All data measured are represented as means ± SEM with dots representing n per group using two-way ANOVA followed by Tukey’s multiple comparisons test. **(J-K)** Pearson’s correlation calculated between the mRNA levels of *Isg15* and fatty acid enzymes, *Acat1* **(J)** and *Cpt1a* **(K).** Asterisks represent \**p* < 0.05; \*\**p* < 0.01; \*\*\**p*< 0.001. n.s. - not significant.

### ISG15 functions as a negative FAO molecular switch during type 1 IFN signaling

Given these inverse correlations between FAO enzyme and *Isg15* transcript levels, and prior data showing that ISG15, as an ubiquitin-like molecule ISGylates metabolic enzymes to modify protein stability, dimerization and activity (17, 22, 23) we employed CRISPR-Cas9 to deplete *Isg15* in BMDMs. The extent of *Isg15*depletion was > 95% (Supplemental Figure 5A-B). We then employed mass spectroscopy based proteomics to quantify the relative levels of FAO enzymes comparing DMXAA treated control and *Isg15*-depleted BMDMs (IKO). IKO BMDMs showed increased levels of CPT1a, HADHA, HADHB, long chain and medium chain acyl-CoA dehydrogenase (ACADVL and ACADM) and ACAT1 compared to controls, following DMXAA stimulation (Figure 5A). Concurrently transcript levels encoding *Cpt1* and *Hadhb* were similarly induced by DMXAA in IKO BMDMs, although this change did not reach significance for *Acat1* (Supplemental Figure 5C-E).) Furthermore, the steady-state protein levels of Cpt1a, Hadha, Hadhb and Acat1, were elevated at baseline in IKO BMDMs and induced to a greater extent than in the control gRNA BMDMs in response to acute DMXAA administration (Figure 5B and Supplementary Fig 5F-I). Additionally, endogenous co-immunoprecipitation in BMDMs showed that antibodies to CPT1a and Acat1 could co-precipitate Isg15 (Figure 5C-D) suggesting ISGylation of these two enzymes. Finally, we validated this interaction between FAO and ISG15 using an integrative bioinformatics approach. Here, comparison of protein levels between WT and IKO BMDMs were integrated with the ISGylome protein dataset obtained post infection with *Listeria monocytogenes.*(*24*) The analysis revealed 211 overlapping proteins, between the DE proteins from our proteomics data compared to the published ISGylome dataset (Figure 5E). Pathway enrichment of the overlapping proteins highlighted pathways associated with fat metabolism constituted 3 of the top 10 pathways affected by ISGylation (Figure 5F).

**Figure 5:**
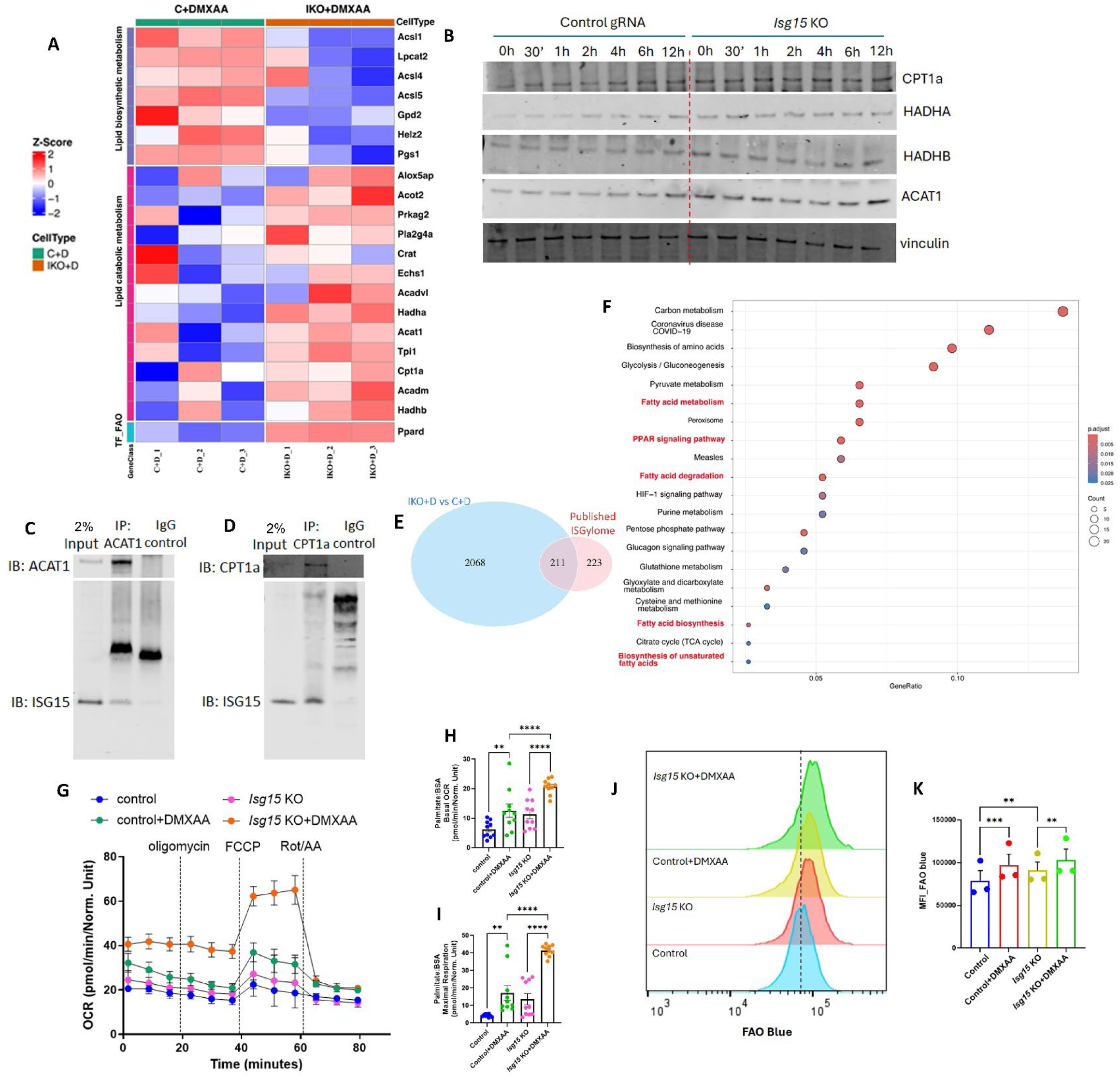
Mitochondrial FAO is negatively regulated by ISG15 during type 1 IFN response. Control gRNA treated and *Isg15* KO BMDMs were treated with 10 μg/mL DMXAA for 4h (n=3 per group) and the cells were subjected to mass spectroscopy based proteomic analysis. **(A)** Heat map representation of levels of proteins associated with fat metabolism as indicated in the sidebars in DMXAA treated control and *Isg15* KO BMDMs. Abbreviations – LB – lipid biosynthesis, LC – Lipid catabolic metabolism, TF – transcription factor. **(B)** Representative immunoblots of key FAO enzymes - CPT1a, HADHA, HADHB, ACAT1 and vinculin in control and *Isg15* KO BMDMs stimulated with DMXAA for indicated timepoints. **(C-D)** Endogenous immunoprecipitation using anti-ACAT1 and anti-CPT1a antibody from lysates of IFN-β (50 ng/mL for 16 h) treated BMDMs, followed by immunoblot analysis using anti-ISG15 antibody. **(C)** Immunoblot representation of endogenous interaction between ACAT1 and ISG15. **(D)** Immunoblot representation of endogenous interaction between CPT1a and ISG15. Two % of the total lysates were used as input controls. **(E)** Venn diagram representation of overlapping proteins from the proteomics data in DMXAA treated control versus *Isg15* KO BMDMs with previously reported ISGylome (PRIDE database identifier no. PXD011513) to identify ISGylated proteins in our proteomics data. The overlapping 211 ISGylated proteins were subjected to pathway enrichment analysis. **(F)** Dot plot of the ClusterProfiler pathway from the 211 ISGylated proteins of a total 2279 DE proteins in DMXAA treated control versus *Isg15* KO BMDMs. The x-axis shows the Gene Ratio, defined as the proportion of input genes (gene names encoding proteins) mapped to a given pathway. Circle size corresponds to the number of genes (Count) involved in each pathway, while the color gradient indicates the adjusted p-value (p.adjust), with red colors reflecting higher statistical significance. Pathways shown represent the top 20 pathways that met the predefined significance threshold.**(G)** Seahorse analysis of OCR in *Isg15* KO and control BMDMs on DMXAA stimulation after sequential introduction of oligomycin, FCCP and Rotenone/Antimycin A (n=10 per group). **(H-I)** Histograms of basal OCR **(H)** and maximal respiration **(I)** of seahorse analysis in G. **(J)** Representative data of flow cytometry based measurement of fluorescence intensity of FaO Blue in control and *Isg15* KO BMDMs with or without DMXAA stimulation. **(K)** Histograms of geometric mean fluorescence intensity of FaO Blue in control versus *Isg15* KO BMDMs with or without DMXAA stimulation. Asterisks represent \*\**p* < 0.01; \*\*\**p*< 0.001.

At a functional level, the rate of FAO in IKO BMDMs was significantly induced as measured by basal and maximal OCR using palmitate as substrate (Figure 5G-I). This was confirmed by FAO blue assessment of endogenous FAO by flow cytometry and microscopy (Figure 5J-K and Supplementary Fig 6A-B). Collectively, these data suggests ISG15 conjugates and modulates the stability of key FAO enzymes by ISGylation to inhibit FAO during heightened type 1 IFN response. Interestingly, the increase in transcript levels encoding the FAO enzymes in the absence of ISG15 implicates additional pre-translational regulatory events in the FAO pathway. Although not the focus of this study, PPARs are well established transcriptional regulators of FAO enzyme encoding genes.(25) In accordance with these previous findings, we showed increased induction of the transcript levels encoding *Ppara* and *Ppard* in IKO BMDMs (Supplementary Figure6 C-D).

### Human clinical studies reflects the interplay of ISG15, FAO and type 1 IFN

We next explored the possibility that ISG15 negatively regulates FAO to control type 1 IFN response. Consistent with our prior data, the type 1 IFN response was enhanced at earlier timepoints in IKO BMDMs in response to DMXAA stimulation measured by activation of STAT1, STAT2 and IRF3 in IKO BMDMs (Figure 6A). We then employed in-silico approaches to assess whether this same regulatory interactions were evident in human studies. Gene array analysis from INFα-stimulated PBMCs from ISG15-deficient patients (GSE60359) exhibited an augmented type 1 IFN response genes induction compared to healthy individuals (Figure 6B).(26) We then assayed these same transcript effects in human patients with known type 1 IFN-mediated diseases including discoid lupus erythematous (DLE) and systemic lupus erythematous (SLE) (GSE179633).(27) Here, analysis of single cell RNA sequencing from dermal skin tissues of healthy volunteers, DLE and SLE patients identified different immune cell types as represented in the respective UMAPs (Supplementary Figure 7G-I). Dot plot analysis revealing the expression pattern of FAO enzymes across the different immune cell type, suggest marked reduction of genes of FAO enzymes in monocytes, macrophages and dendritic cells in both DLE and SLE samples compared to healthy counterpart (Figure 6C-E). Additionally, DLE and SLE showed significant increase in the expression of STAT1, STAT2 and ISG15 (Figure 6C-E). We next isolated CD14^+^ myeloid cells from PBMCs of SLE patients and healthy volunteers (Figure 6F): cells from 9 SLE participants and 9 age and sex-matched controls were studied. Flow cytometric measurement of FAOBlue fluorescence showed decreased FAO in SLE compared to healthy volunteers (Figure 6G). The decrease in FAO rate was accompanied with decreased gene expression of FAO enzymes and increased transcript levels of ISG15 (Figure 6I-M). Interestingly, this negative effect of ISG15 on FAO enzymes was also evident in the microarray data ( GSE60359),(28) in PBMCs from ISG15 deficient patients. ISG15 deficient patients showed elevated expression of key FAO enzymes- *CPT1A*, *ACAT1* and *HADHA* (Figure 6N).

**Figure 6:**
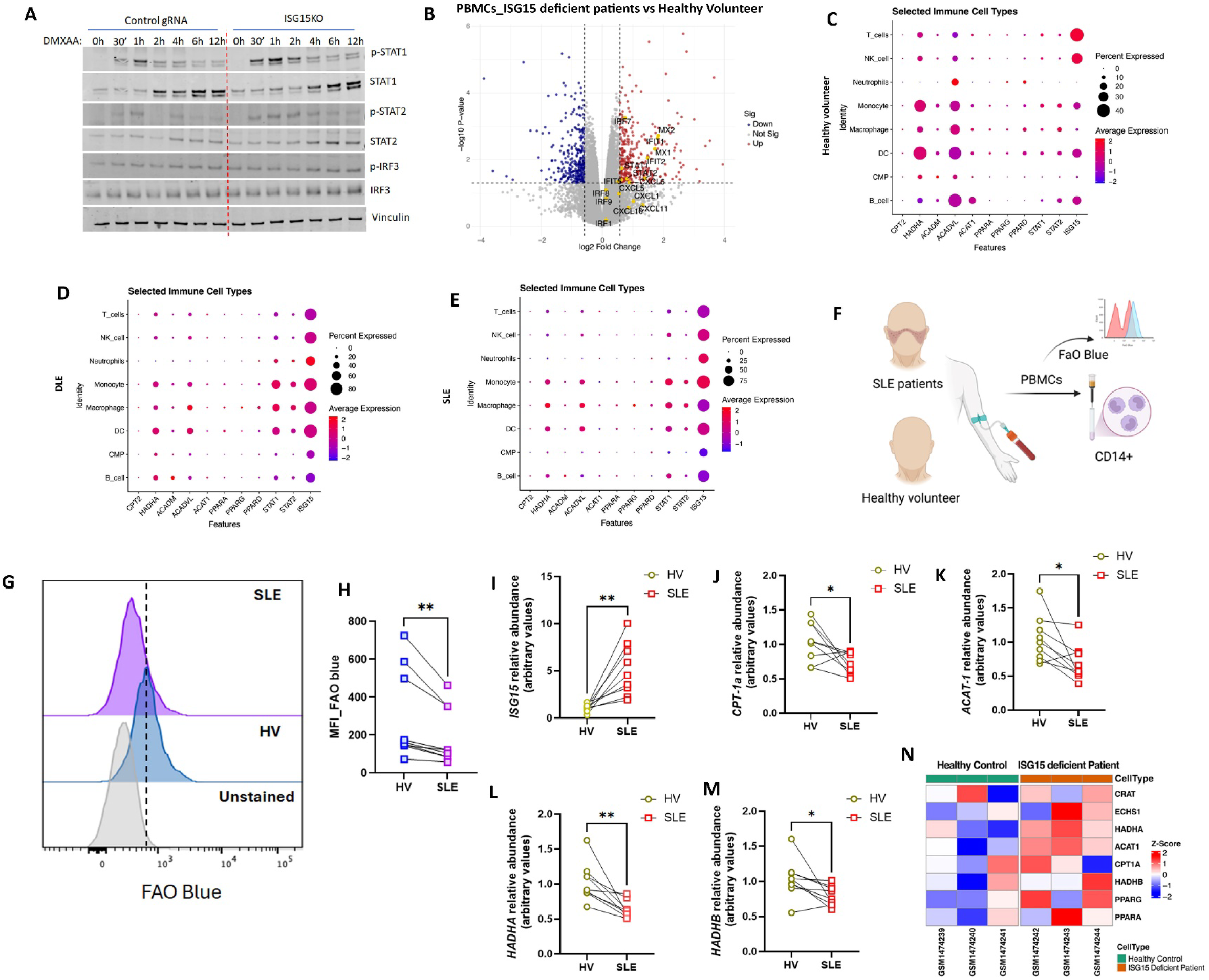
ISG15 mediated feedback inhibition of FAO is reflected in lupus patients. **(A**) Representative immunoblots of p-STAT1, STAT1, p-STAT2, STAT2, p-IRF3, IRF3 and vinculin from control and *Isg15* KO BMDMs stimulated with DMXAA for different timepoints (0-12h).(n=5) **(B)** Volcano plot representation of differentially expressed genes in IFN-α treated PBMCs from ISG15 deficient patients compared to IFN-α treated PBMCs from healthy volunteers (GSE60359). x-axis represents log2 fold change in gene expression, and y-axis represents the −log10 adjusted p-value. Each point corresponds to a gene. Genes with large fold changes and high statistical significance appear toward the top left and top right of the plot, highlighting significantly downregulated and upregulated genes, respectively. Genes associated with type 1 IFN response has been highlighted as yellow dots. **(C-E)** Dot plot representation of scRNA analysis to understand the expression pattern of FAO associated genes in different immune cells from dermal tissues of healthy volunteers **(C)**, DLE **(D)** and SLE **(E)** patients. Dot size represents the percentage of immune cells expressing each gene and dot color represents the average expression level of indicated genes. **(F)** Schematic of the clinical validation study to determine FAO in CD14+ myeloid cells from PBMCs of control vs. SLE patients. PBMCs isolated from blood of 9 SLE patients and age matched healthy volunteers were incubated with 5 µM FaO Blue for 1h followed by staining with APC conjugated anti CD14 antibody. The CD14+ populations were gated and analyzed for fluorescent intensity of FaO blue by flow cytometry. **(G)** Representative data of flow cytometry based measurement of FaO Blue intensity in CD14+ gated PBMCs from SLE patient and respective age matched healthy volunteer. **(H)** Dot plot of paired geometric mean fluorescent intensity of FaO Blue in CD14+ gated PBMCs from SLE patient and respective age matched healthy volunteer (n=9). **(I-M)** Quantitative RT-PCR analysis of *ISG15* **(I)**, *CPT1A* **(J)**, *ACAT1* **(K)**, *HADHA* **(L)** and *HADHB* **(M)** in CD14+ myeloid cells isolated from PBMCs of SLE patients and respective age matched healthy volunteers. Data were normalized to 18*S* rRNA. Paired t-test. **(N)** The heatmap shows expression profiling by array data from the GSE60359 dataset, comparing healthy controls and ISG15-deficient patients. Rows represent the genes of key FAO enzymes *CRAT*, *ECHS1*, *HADHA*, *ACAT1*, *CPT1A*, *HADHB*, *PPARG*, and *PPARA*, while columns represent six individual samples. Expression values were normalized using quantile normalization and scaled to z-scores. Colors indicate relative gene expression levels, as shown in the color bar. Asterisks represent - \**p*< 0.05; \*\**p* < 0.01; \*\*\**p*< 0.001. n.s. - not significant.

## Discussion

In this study, we show a dynamic and biphasic relationship between fatty acid oxidation and type I interferon signaling. Moreover, we show that the type I interferon responsive gene, ISG15 plays a central post-translational negative feedback regulatory role in blunting fatty acid oxidation to dampen subsequent type I interferon signaling.

More broadly, in this study we investigated the interaction between mitochondrial fatty acid oxidation (FAO) and the type I IFN response. We demonstrate that activation of the STING pathway, a key inducer of type 1 IFN, enhanced mitochondrial oxygen consumption in myeloid cells when palmitate was included as a oxidative substrate. This metabolic shift suggested a reprogramming towards increased mitochondrial FAO, which was further supported by the upregulation of FAO-associated enzymes. Disruption of FAO through genetic depletion of *Cpt1a* resulted in reduced expression of type 1 IFN-stimulated genes following STING agonist or LPS stimulation. In *Cpt1a* -deficient BMDMs, we observed decreased levels of STAT1, STAT2, and IRF3, which correlated with reduced histone H3 acetylation at lysines 9 and 14. These findings support an FAO-dependent epigenetic mechanism regulating type 1 IFN responses, a regulatory component that aligned with our prior findings in *Acat1*-deficient murine macrophages.(7)

Using an *in vivo* LCMV infection model, we then confirmed that type 1 IFN induction, in an apparent feed-forward loop, augmented FAO in myeloid cells at early time points (days 1 and 3) post-viral infection. These data directly correlating the rates of FAO with type I interferon signaling are consistent with our prior in *Acat1* knockout data in monocytes.(7) Additionally, pharmacological inhibition of FAO using high doses of etomoxir in plasmacytoid dendritic cells reduced the type 1 IFN responses in response to LCMV infection.(15) Interestingly, FAO levels declined by day 5 post infection, implicating a temporal regulatory mechanism exerting feedback inhibition of mitochondrial FAO. This reduction in FAO correlated with the reduction of transcript levels encoding interferon regulatory proteins, with the exception of ISG15, where transcript levels were elevated. The dynamic interaction between the type 1 IFN and mitochondrial oxidative phosphorylation, has also been shown where the dosing of IFN-β elicits a biphasic effect on macrophage mitochondrial oxygen consumption rates, although this was in the presence of glucose and glutamine and without fatty acid substrates.(19) Additionally, the role of fatty acid oxidation in response to different fatty acid species have begun to be identified in other inflammatory programs, including in the NLRP3 inflammasome in resident skin macrophages.(29) Together these data suggest that the control of FAO may have a broader dynamic and temporal role in immunometabolism and immunoregulation.

The interferon responsive gene ISG15, is a ubiquitin-like modifier, and strongly induced by type 1 IFN. It functions either through ISGylation of target proteins to influence their activity and/or stability or, in its unconjugated form, functions as an inflammatory cytokine.(30) Its direct immunomodulatory effect is evident where ISG15 confers negative feedback on type 1 IFN signaling by conjugating to RIG-I to diminish its level and blunt IFN promoter activity.(31) Additionally ISG15 non-covalently interacts with Ubiquitin-Specific Protease 18 (USP18) to prevent its degradation and thereby enhances the USP18 inhibitory effects on type 1 IFN signaling.(32) Moreover, ISG15 inhibits type 1 IFN mediated JNK activation by conjugating with filamin B to impede the filaminB:RAC1 interaction. This dissociation, in turn, attenuates type 1 IFN-induced JNK signalling.(33) Another mechanism whereby ISG15 can modulate immune function is via autophagy, which itself is linked to type 1 IFN signaling,(34) and here ISGylation modifies autophagy mediators and increases autophagic flux.(24) Beyond these established antiviral roles, ISG15 has recently been implicated in host metabolic regulation, where ISG15 levels modulate levels of glucose, fat oxidation and lipid turnover proteins.(17, 35, 36) Furthermore, in other tissues exposed to infectious trigger pathways, ISG15 is shown to induce hepatic gluconeogenesis,(22) and to blunt glycolysis in adipose tissue to modulate thermogenesis.(35) This targeting of metabolism may be context specific as exogenous ISG15 has been shown to promote glycolysis and lactate production to enhance fibroblast migration in the tumor microenvironment.(37) To date, the mechanisms whereby ISG15 mediated changes in metabolic enzyme levels modulate immunity is less well characterized. Here we show that ISG15 directly interacts with key FAO enzymes, including CPT1a and ACAT1, aligning with prior ISGylome datasets identifying these enzymes as ISG15 targets.(17, 24, 35) Since it is challenging to pull down ISG15 using antibody,(38) most published ISGylome study in different cell types have incorporated tagged based pull down strategies to identify ISGylated proteins. Using these indirect approaches, FAO enzymes have been identified in several ISGylome datasets. In this study we employed the more direct approach, i.e. without using tagged ISG15, where FAO enzymes were used for immunoprecipitation followed by the assessment of endogenous ISG15 by immunoblot analysis. This approach showed the interaction between ISG15 and CPT1a and ACAT1. Furthermore we identify ISG15 as a negative molecular switch for mitochondrial FAO during a heightened or prolonged type 1 IFN response. Additionally, the proteomic data in the absence of ISG15 and interaction studies with published ISGylome datasets support that this regulation functions at the level of FAO enzyme ISGylation and the reduction in FAO metabolism. Together, these data are expanding our understanding of the interplay between FAO and type 1 IFN regulation. Interestingly, humans with ISG15 deficiency show a heightened type 1 IFN response (26) and our in-silico analysis from these individuals show the association between ISG15 deficiency and increased transcript levels encoding FAO enzymes.

Nevertheless, an additional level of regulation may be operational in linking to ISG15 mitochondrial FAO. This was initially suggested where, in response to Vaccinia viral infection, ISG15 knockout BMDMs showed increased expression of the transcripts encoding peroxisome proliferator activated receptors gamma (PPARγ) and of the PPARγ coactivator-1a (PGC-1α).(17) PGC-1α and PPARγ are well established as essential transcriptional machinery in driving mitochondrial FAO.(39) As shown in our supplemental data, different PPAR isoforms, i.e. PPARα and PPARδ were also upregulated in ISG15 knockout BMDMs. These isoforms similarly are pivotal for FAO in different tissue types.(40, 41) Although we did not explore this mechanism further, these data add yet another level of regulation to the complex array of counter-regulatory control nodes between FAO and type 1 IFN signaling.

Finally, our findings in primary murine BMDMs appear to align with expression profiling studies in humans with known interferonopathies. This is evident in both DLE and SLE where FAO is reduced and ISG15 expression is elevated. Single-cell RNA sequencing data from DLE and SLE patients confirm reduced FAO enzyme levels alongside increased ISG15 expression, reinforcing the role of ISG15 as a suppressor of mitochondrial fat oxidation during excessive type 1 IFN activity.

Together these findings strongly support a novel regulatory axis whereby ISG15 restricts type 1 IFN responses through inhibition of FAO. Moreoever, our findings reveal a complex multidimensional regulatory network linking fat metabolism and type I interferon signaling. Further studies will be required to dissect out the numerous epigenetic, transcriptional, and post-translational control nodes in these interactions between FAO, type I interferon and ISG15. Interestingly, obesity is an independent risk factor for excessive inflammation and worse functional capacity in SLE.(42, 43) The link between obesity, increased fat oxidation and elevated type I interferon has also been implicated in a small human cohort.(7) Together, all of these findings point to the importance of mitochondrial fat oxidation in type I interferon signaling and underscore that understanding this regulation is now an integral component of the broader role of metabolism in immune regulation.

## Experimental Methods

### Animal studies and LCMV infection

Wild type C57BL/6 mice (stock no. 000664; Jackson Laboratory) were maintained under specific-pathogen-free facility of National Heart Lung Blood Institute (NHLBI) and provided with food and water ad libitum. Male mice (6-8 weeks of age) were sacrificed for bone marrow derived macrophage (BMDM) studies.

For LCMV infection, 6 weeks old male mice (n=5 per time point) received 2×10^5^ LCMV Armstrong via intraperitoneal injection for different time points (1-8 days). Schematic of the study design is shown in Figure 3A. At the predetermined timepoints post-infection, mice were euthanized and splenectomized for splenocyte analysis. Animal experiments were approved by the NHLBI Animal Care and Use Committee.

### Isolation of BMDMs and cell culture experiments

Bone marrows were extracted from C57BL/6 mice femurs and plated in DMEM (10% FBS) along with 25 ng/mL murine M-CSF ( at 37^0^ C and 5% CO_2_ for 5 days to differentiate into BMDMs. The M-CSF media were aspirated and fresh DMEM (10% FBS) without M-CSF were added for 24 h. 3 × 10^5^ BMDMs were plated per well in a 24 well plate and were treated with 10 μg/mL 5,6-dimethylxanthenone-4-acetic acid (DMXAA) or 50 ng/mL recombinant IFN-β or 10ng/mL lipopolysaccharide (LPS) for indicated time points. The supernatant were collected from DMXAA and LPS treated cells and centrifuged to remove cells and debris and stored at –80^0^ C for analysis of IFN-β cytokine levels by ELISA (R&D Systems), while the cells were washed with phosphate buffer saline (PBS) post treatment and RNA or protein were isolated.

### CRISPR Cas9 mediated gene editing

Alt-R CRISPR-Cas9 system (IDT) was used to knockout CPT1a and ISG15 from primary BMDMs.(44) sgRNAs were made by incubating either *Cpt1a* or *Isg15* crRNA along with tracr RNA (95^0^ C −1 min, 60^0^ C- 1 min and hold at 25^0^ C). Following which the CRISPR ribonucleoprotein (RNP) complexes were constructed by incubating the sgRNA along with Cas 9 for 15 min at 37^0^ C followed by a hold at 8^0^ C. 10 × 10^6^ BMs from WT C57Bl/6 mice were washed twice with PBS and dissolved in P3 Primary Cell Buffer and transferred to nucleofection chamber. CRISPR RNP complexes were delivered to the BMDMs by nucleofection using the CM137 pulse program on Amaxa 4D-Nucleofector (Lonza). Pre-warmed media was immediately added to the nucleofected cells and incubated at 37^0^ C for 30 min before seeding on plates containing BMDM differentiation media. Knock out was confirmed by immunoblot analysis.

### Metabolic assays

Real-time measurement of oxygen consumption rates (OCR) of BMDMs treated with DMXAA or IFN-β for indicated times were performed on a XFe-96 Seahorse Extracellular Flux 159 Analyzer (Agilent), using the Mitostress kit following instructions from the manufacturer. OCR was measured in XF DMEM media (containing 10mM glucose, 2mM L-glutamine, and 1mM pyruvate) after sequential injections of 2 µM oligomycin, 1.5 µM FCCP (carbonyl cyanide-4-(trifluoromethoxy)-phenylhydrazone), and 0.5 µM antimycin A plus rotenone. In order to access the relative oxygen consumption from fat oxidation DMXAA stimulated BMDMS were pre-treated for 15 min with etomoxir (10 µM) concentration. Acute changes of etomoxir in maximal respiration of IFN-β stimulated cells was assessed by injecting 20µM etomoxir into the wells after FCCP injection.

To evaluate the capacity of BMDMs to oxidize external fatty acids, the cells were plated in 96 well Seahorse plate and incubated overnight in substrate-limited media (XF DMEM media containing 0.5 mM glucose, 1mM glutamine, 1% FBS and 0.75mM L-carnithine) along with DMXAA or IFN-β treatment for indicated timepoints. The following day and after respective treatments, the cells were washed with PBS and switched to FAO assay medium (XF DMEM media containing 111 mM NaCl, 4.7 mM KCl, 1.25 mM CaCl₂, 2.0 mM MgSO₄, 1.2 mM Na₂HPO₄, 0.5 mM glucose, 0.75 mM carnitine, and 5 mM HEPES) for 1 hour. After a 15-minute pretreatment with etomoxir (10 μM), when indicated, palmitate–BSA (200 μM palmitate conjugated to 34 μM BSA) was added, and OCR was recorded following sequential injections of 2 µM oligomycin, 1.5 µM FCCP and 0.5 µM antimycin A plus rotenone with or without 15 min of pre-treatment with etomoxir (10 µM) concentration. OCR values were normalized to number of cells.

Splenocytes from LCMV infected cells were collected and plated in Seahorse XF RPMI Base media (pH=7.4, without phenol red) along with 5 μM FaO Blue as the fatty acid source for 60 min. Post 60 min of incubation with FaO blue the splenocytes were washed and stained with PE-Rat anti-CD11b antibody in a 1:100 ratio for 20 min in dark. The splenocytes were then washed with PBS+1%FBS and flowcytometric analysis performed immediately. CD11b+ cells were isolated from the splenocytes using positive magnetic selection and isolated CD11b+ cells were seeded along with 5 μM FaO blue in XF RPMI base media for 60 min in poly-L-lysine (0.1%) coated 8 well chambered slides or subjected to RNA isolation. Post 60 min of incubation with FaO blue the microscopic slides were washed with PBS and fixed using 2% para-formaldehyde.

### RNA Isolation and Quantitative PCR (qRT-PCR) analysis

BMDMs were treated with DMXAA for indicated time points. Following which the cells were washed with PBS and total RNA was isolated using the NucleoSpin RNA kit (Macherey-Nagel), and cDNA was synthesized with the SuperScript III First-Strand Synthesis System for RT-PCR (Thermo Fisher Scientific). For *in vivo* LCMV infection study CD11b+ cells were positively isolated from the splenocytes of infected mice prior suspension in RA1 buffer after indicated time points of infection, while CD14+ cells from the PBMCs of SLE patients were used Quantitative real-time PCR was carried out using FastStart Universal SYBR Green Master (Roche) on a LightCycler 96 System (Roche). Relative gene expression levels were determined by normalizing the cycle threshold (Ct) values to 18S rRNA and analyzed using the 2^–ΔΔCt method. The primer sequences are provided in Supplementary Tables 1 and 2.

### Western Blot analysis

Total cell lysates from BMDMs post treatment were made using radioimmunoprecipitation assay (RIPA) buffer with protease inhibitor cocktail (Roche) and phosphatase inhibitors (Sigma-Aldrich). Proteins were separated on 4-12% NuPAGE Bis-Tris gels (Thermo Fisher Scientific) and transferred to nitrocellulose membrane using the the Trans-Blot Turbo Transfer System (Bio-Rad Laboratories) following the manufacturer’s protocol. Membranes were blocked with 50% Odyssey Blocking Buffer in PBS-T [PBS containing 0.1% Tween-20] and incubated overnight at 4 °C with the appropriate primary antibodies. The following antibodies were used: STAT1, phospho-STAT1, STAT2, phospho-IRF3, ISG15, acetyl-Histone H3(Lys9/Lys14), Histone H3 (Cell Signaling Technology); phospho-STAT2 (EMD Millipore); IRF3 (Abcam); Cpt1a, HADHB (proteintech); HADHA (Invitrogen); ACAT1 and vinculin (Sigma-Aldrich). After primary incubation, membranes were incubated with IRDye secondary antibodies for 1 hour at room temperature. Blots were visualized using the Odyssey CLx Imaging System (LI-COR Biosciences). Protein band intensities were quantified with ImageJ software and normalized to vinculin.

### Endogenous co-immunoprecipitation

30 × 10^6^ BMDMs were treated with 50ng/mL IFN-β for 16h followed by washing and lysis in Pierce IP lysis buffer. The cells were lysed by sonication and centrifuged at 13000g for 10 min for removal of debris. The supernatant was collected and protein concentration estimated. 2% of the lysate were removed and set aside for input. The lysates were kept overnight in rotation at 4^0^ C along with protein G beads priorly incubated with either mouse anti-CPT1a or goat anti-ACAT1 antibodies. Post overnight incubation, the lysates were removed by magnetic separation followed by washing the beads and eluting in SDS-lameli buffer containing 2% β-mercaptoethanol. Immunoprecipitation was confirmed using rabbit anti-CPT1a or rabbit anti-ACAT1 antibodies. Co-immunoprecipitation of both proteins with ISG15 was confirmed using anti ISG15 antibody. Anti-mouse IgG or anti-goat IgG were used as respective controls.

### Liquid chromatography-mass spectroscopy sample preparation

*Isg15* KO BMDMs treated with DMXAA were washed with PBS and then lysed on ice in 100 µL EasyPep™ lysis buffer supplemented with protease inhibitors (Roche), resuspended by gentle pipetting, flash-frozen on dry ice, thawed, and centrifuged at 14,000 rpm for 10 min at 4 °C, after which the supernatant was collected. Proteins were reduced and alkylated by adding 4 µL 500 mM chloroacetamide and 1 µL 500 mM TCEP to 100 µL lysate, denatured at 95 °C for 10 min, cooled, and digested overnight at 37 °C with 10 µL trypsin (0.1 µg/µL in 50 mM HEPES, pH 8.0). Peptides were quantified using a Thermo colorimetric assay at 480 nm with a BSA standard curve. For TMT labeling, 10 µg peptide per sample was adjusted to 100 µL HEPES buffer, mixed with TMT reagent reconstituted in 40 µL acetonitrile, incubated ≥1 h at room temperature, and quenched with a solution of 5% hydroxylamine and 20% formic acid before samples were combined and cleaned using the EasyPep™ MS Sample Prep Kit. Dried peptides were resuspended in 50 µL 0.1% TFA, vortexed for 15 min, centrifuged at 10,000 rpm for 3 min, transferred to autosampler vials, and 10 µL was injected for LC–MS analysis.

### Proteomics data analysis

Total proteome analysis was conducted on murine BMDMs samples across four experimental groups (control, Control+DMXAA, *Isg15* KO, and *Isg15* KO+DMXAA; n = 3 per group, 12 biological replicates total), resulting in quantification of 3,018 expressed proteins. To reduce technical variability, protein intensities were normalized by total intensity normalization, whereby the summed protein intensity of each sample was scaled relative to the sample with the highest total intensity to generate a normalized expression matrix. Differential protein expression was assessed using the limma package (v3.66.0) in R (v4.5.1), with pairwise comparisons performed between experimental groups and multiple testing correction applied using the false discovery rate (FDR) method; proteins with an adjusted p-value < 0.05 and an absolute log₂ fold change > 0.58 were considered significantly differentially expressed. Data visualization, including volcano plots, expression distributions, and heatmaps, was performed using ggplot2 (v4.0.1) and pheatmap (v1.0.13), while overlap analyses between differentially expressed protein sets and public gene sets were conducted using ggvenn (v0.1.19) and VennDiagram (v1.7.3). Functional enrichment analysis was carried out using clusterProfiler (v4.18.2), with gene annotation and identifier mapping based on the org.Mm.eg.db database (v3.22.0), and pathway enrichment was evaluated using KEGG analysis to identify significantly perturbed biological processes and signaling pathways associated with the observed proteomic changes.

### Human Studies

Healthy volunteers and systemic lupus erythematous participants were recruited onto a National Heart and Lung Institute natural history protocol ([NCT01143454](https://clinicalstudies.info.nih.gov/ProtocolDetails.aspx?id=2010-H-0126)). Nine female SLE and age-matched (2.7±0.6 years) healthy volunteers (with ages ranging from 36 to 62 years), signed an informed consent prior to donating blood for analysis for this study. The SLE participants were stable on steroids, ± methotrexate and/or ± hydroxychloroquite, and no SLE individuals were on biologic therapeutics. Primary human peripheral blood mononuclear cells (PBMCs) were isolated from whole blood by density gradient centrifugation using Lymphocyte Separation Medium (MP Biomedicals). Part of the isolated total PBMCs from both groups were plated onto 12 well plate in XF RPMI media (without phenol red) along with 5 μM FaO blue for 60 min at 37^0^ C. Post 60 min the cells were washed with PBS and stained with APC-Mouse anti human CD14+ in dark for 20 mins. Flow cytometric analysis of FAO was done immediately after washing the cells in PBS+1% FBS. Monocytes were subsequently purified from PBMCs by negative selection using the CD14+ Monocyte Isolation Kit (Miltenyi Biotec). 2.5 × 10^6^ monocytes were plated on per well onto a 12 well plate for assessment of FAO using FaO blue in XF RPMI media, as described previously or total RNA isolation.

### scRNA sequencing analysis

Single-cell RNA sequencing data were obtained from the publicly available dataset GSE179633, from which dermal tissue samples were selected, comprising 16 samples in total (4 healthy controls, 5 discoid lupus erythematosus, and 7 systemic lupus erythematosus). Data were analyzed using Seurat (v5.4.0) in R, with individual samples imported and merged into a single Seurat object followed by standard quality control filtering based on gene counts and mitochondrial gene expression, log-normalization using the LogNormalize method (scaling to 10,000 counts per cell and log transformation), and data scaling. Cell-type annotation was performed using SingleR (v2.12.0) with reference datasets from the celldex package (v1.20.0) to enable automated and consistent cell labeling. Differential gene expression analyses comparing SLE versus healthy controls and DLE versus healthy controls were conducted using the Wilcoxon rank-sum test implemented in Seurat’s FindMarkers function, with multiple testing correction applied using the Benjamini–Hochberg false discovery rate; genes with an adjusted P value < 0.05 were considered significantly differentially expressed. Visualization of cell-type distributions and differential expression results was performed using Seurat visualization tools and ggplot2, enabling assessment of transcriptional heterogeneity across disease and control conditions.

### Statistical analysis

Statistical analyses were conducted using GraphPad Prism 10. Data are presented as mean ± SEM unless otherwise specified. For the *in vivo* and *in vitro* studies, *n* represents the number of biological replicates per group and is reported in the figure legends. Comparisons between two groups were performed using paired or unpaired two-tailed Student’s *t* tests. Comparisons among more than two groups were analyzed using one-way analysis of variance (ANOVA) followed by appropriate multiple-comparison tests (Dunnett’s or Tukey’s). Two-way ANOVA was applied when two independent variables were present. A *P* value < 0.05 was considered statistically significant.

## Author Contributions

Conceptualization: A.K.G. and M.N.S. Funding acquisition: W.J.L., C.K. and M.N.S. Investigation: A.K.G., J.W., P.D., S.D., R.S., K.H., C.C., and E.E.W. Methodology: A.K.G., J.W., P.D. and S.D. Validation: A.K.G., J.W., K.H. and R.J.K. Supervision: A.K.G., J.W., R.D.H. and M.N.S. Clinical Study Support: R.D.H. and M.N.S. Formal analysis: A.K.G., J.W., P.D., K.H. and S.D. Project administration: A.K.G and M.N.S. Visualization: A.K.G. and P.D. Software: A.K. G. and P.D. Writing—original draft: A.K.G., and M.N.S. Writing—review and editing: A.K.G., J.W., K.H., R.J.K, W.J.L., C.K., E.E.W. and M.N.S.

## Acknowledgements

This research was supported by the Intramural Research Program of the National Institutes of Health (NIH). The contributions of the NIH author(s) were made as part of their official duties as NIH federal employees, are in compliance with agency policy requirements, and are considered Works of the United States Government. However, the findings and conclusions presented in this paper are those of the author(s) and do not necessarily reflect the views of the NIH or the US Department of Health and Human Services. Funding for this study was supported by the NIH intramural programs to MNS (grant HL-005102).

## Notes

### Competing Interest Statement

The authors have declared no competing interest.

