## Supplemental Figs and Tables for "ISG15 orchestrates dynamic crosstalk between mitochondrial fat oxidation and type 1 interferon in myeloid cells"

**Supplementary Data:**

Supplementary Figures – 7

Supplementary Tables – 2

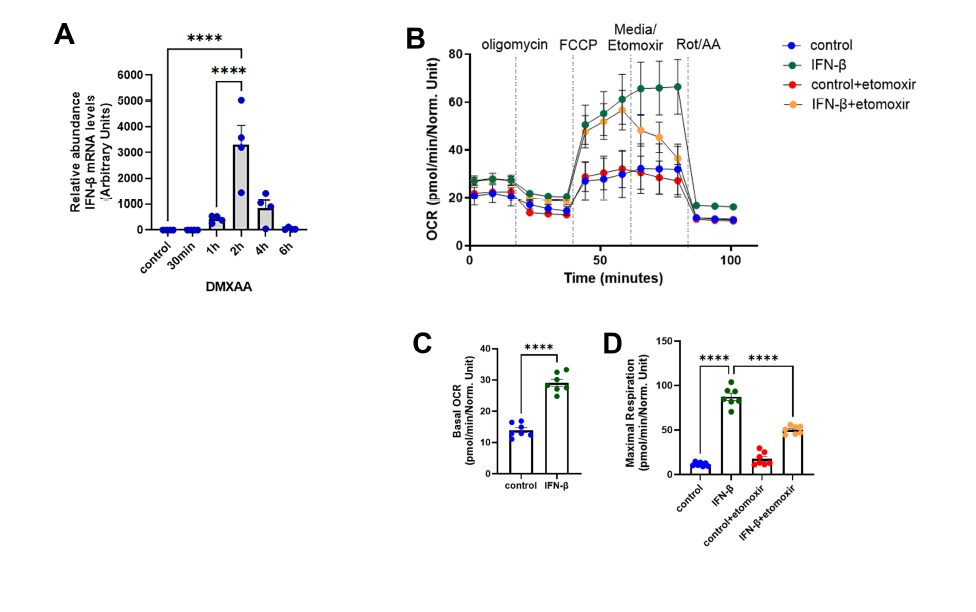

**Supplementary Figure 1: Induction of mitochondrial FAO in myeloid cells by T1IFN**

**(A)** Quantitative RT-PCR analysis of *Ifnb* levels in BMDMs treated with 10µg/mL DMXAA for indicated time points (0-6h) (n=4 per group). Data were normalized to 18SrRNA levels. **(B)** Seahorse analysis of oxygen consumption rate (OCR) of BMDMs stimulated with 50ng/mL IFN-β for 4 h in response to sequential treatment of oligomycin, FCCP, media or etomoxir and Rotenone/Antimycin A (n=7 per group). **(C-D)** Histograms of basal OCR (D) and maximal respiration (E) of Seahorse analysis in B. Results measured were normalized to the cell counts in each corresponding wells. Two-way ANOVA followed by Tukey’s multiple comparisons test. *****p* < 0.0001.

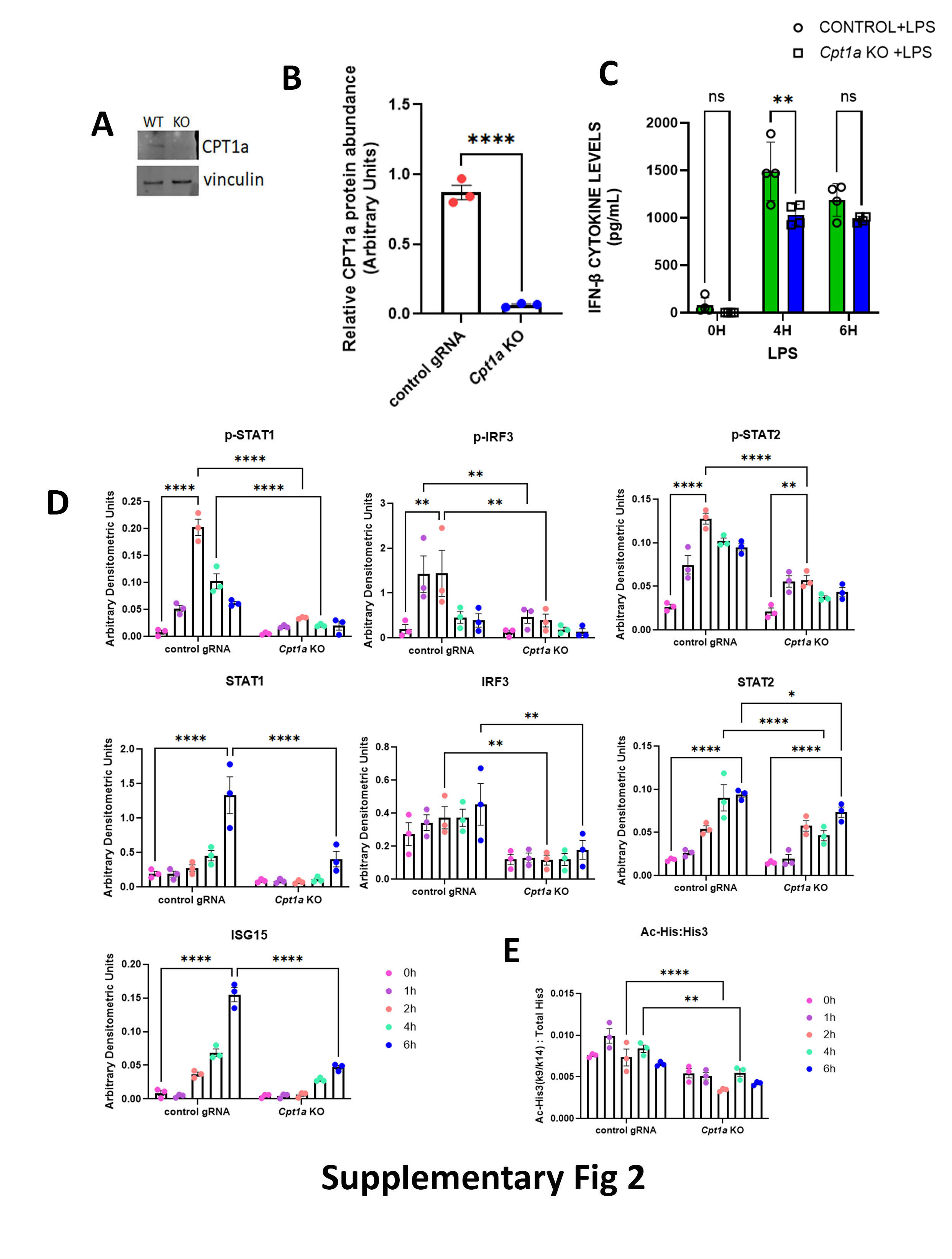

**Supplementary Figure 2: Genetic depletion of *Cpt1a* blunts T1IFN response**

**(A**) Representative immunoblots of CPT1a levels in control gRNA treated and *Cpt1a* KO BMDMs. **(B)** Graphical representation of CPT1a protein levels in control versus *Cpt1a* KO BMDMs (n=3 per group). **(C)** Graphical representation of IFN-β cytokine levels in LPS treated control and *Cpt1a* KO BMDMs (n=4 per group). **(D)** Densitometric analysis of immunoblots of p-STAT1, STAT1, p-IRF3, IRF3, p-STAT2, STAT2 and ISG15 relative to vinculin (n=3 per group). **(E)** Graphical representation of Ac-His3 K9/K14: total His 3, analyzed post densitometric analysis of immunoblots of Ac-His3 K9/K14 and total His3 relative to vinculin (n=3 per group). Two-way ANOVA followed by Tukey’s multiple comparisons test. **P* < 0.05; ***P* < 0.01; ****P* < 0.001. n.s., not significant.

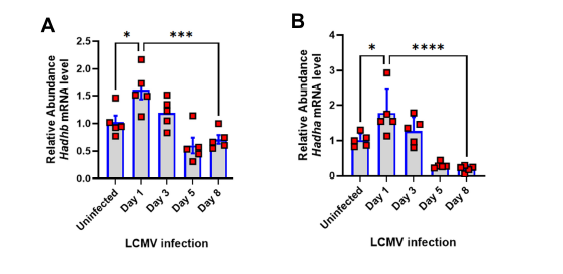

**Supplementary Figure 3: Level of key FAO enzymes in CD11b+ myeloid cells from splenocytes of LCMV infected mice**

**(A-B)** Quantitative RT-PCR of *Hadhb* **(A)** and *Hadha* **(B)** in CD11b+ myeloid cells from splenocytes of LCMV infected mice at indicated points of infection (n=5 per group). Data were normalized to 18S rRNA levels. Two-way ANOVA followed by Tukey’s multiple comparisons test. **P* < 0.05; ***P* < 0.01; ****P* < 0.001. n.s., not significant.

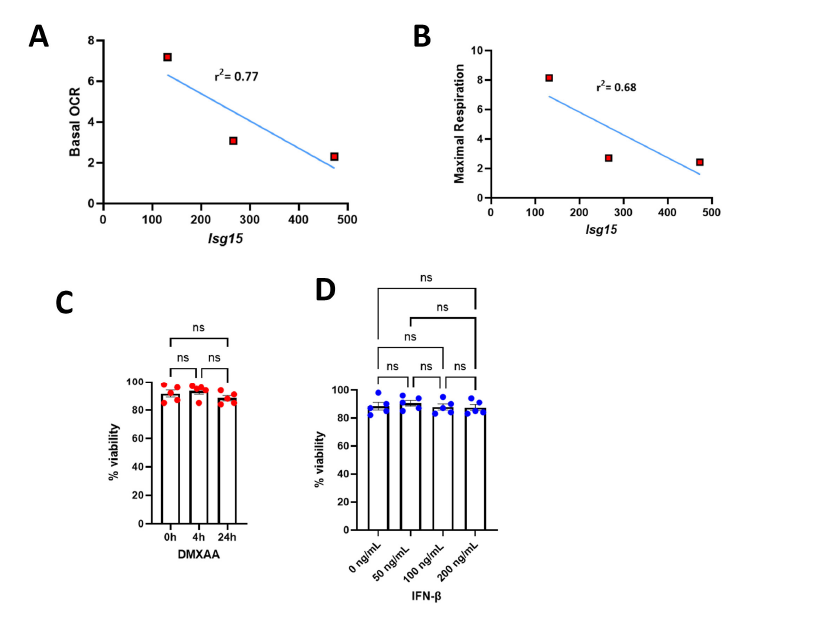

**Supplementary Figure 4: ISG15 negatively correlates with mitochondrial fat oxidation**

**(A-B)** Graphical representation of correlation between *Isg15* levels and basal OCR **(A)** and maximal respiration **(B)** of BMDMs stimulated with increasing concentrations of IFN-β (0ng/mL-200ng/mL) in presence of BSA conjugated palmitate as substrate. **(C-D)** Viability of BMDMs when treated with DMXAA **(C)** for indicated timepoints and increasing concentrations of IFN-β **(D)** as analyzed by trypan blue staining.

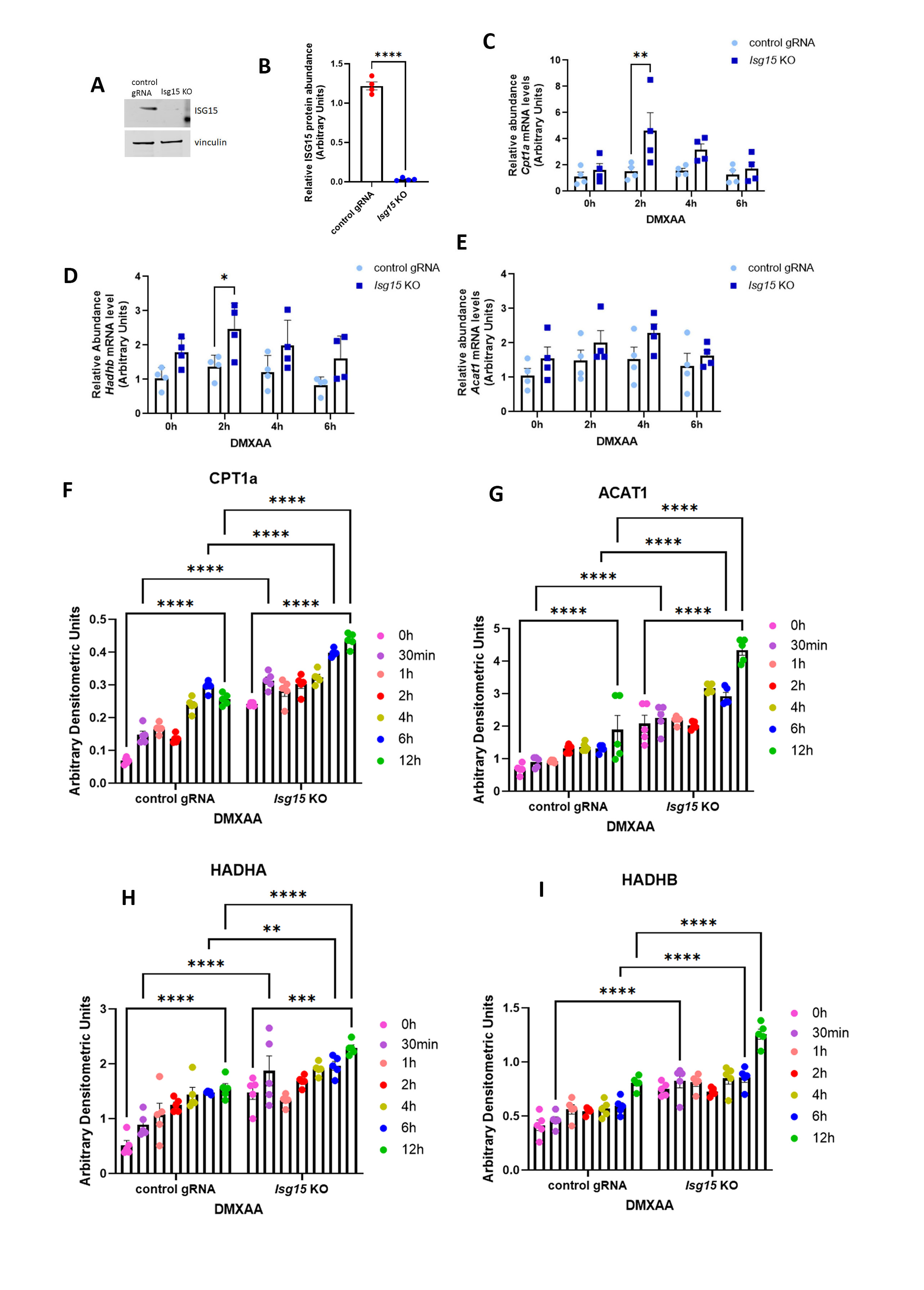

**Supplementary Figure 5: Level of key FAO enzymes in DMXAA treated *Isg15* KO BMDMs**

**(**A) Representative immunoblot of ISG15 levels in control gRNA treated and *ISG15* KO BMDMs. **(B)** Graphical representation of ISG15 protein levels in control versus *Isg15* KO BMDMs (n=4 per group). **(C-E)** Quantitative RT-PCR analysis of key enzymes of FAO-*Cpt1a* **(C)**, *Hadhb* **(D)** and *Acat1* **(E)** in *Isg15* KO BMDMs stimulated with or without 10 μg/mL DMXAA for indicated timepoints (n=4 per group). Data were normalized to 18*S* rRNA and represented as means ± SEM. **(F-I**) Protein quantitation and densitometric analysis of CPT1a **(F)**, ACAT1 **(G)**, HADHA **(H)**, HADHB **(I)** relative to vinculin in DMXAA treated control and *Isg15* KO BMDMs (n=5 per group). Two-way ANOVA followed by Tukey’s multiple comparisons test. **P* < 0.05; ***P* < 0.01; ****P* < 0.001. n.s., not significant.

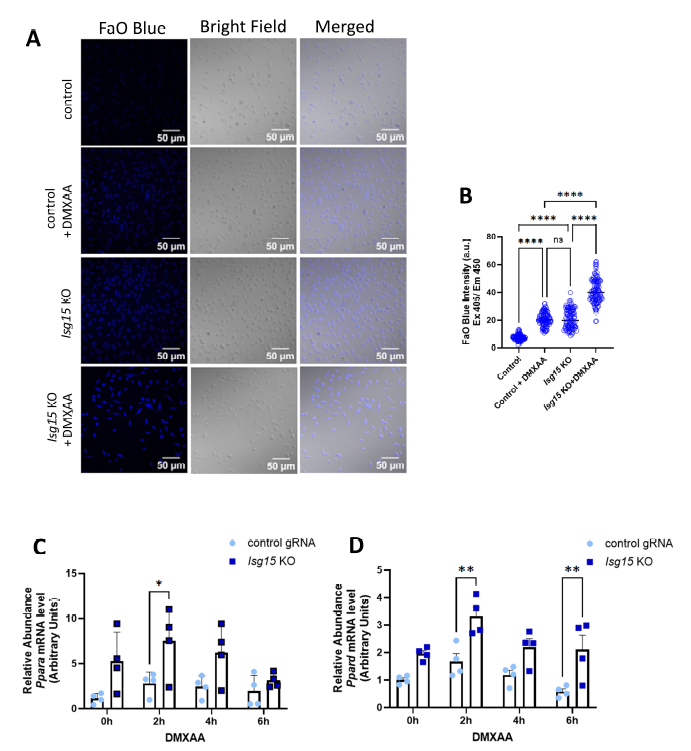

**Supplementary Figure 6: ISG15 mediated negative regulation of mitochondrial fat oxidation**

**(A**) Representative microscopic images of control and *Isg15* KO BMDMs treated with or without DMXAA for 4h followed by incubation with 5 µM FaO Blue. **(B)** Graphical representation of fluorescence intensity of FaO Blue from microscopic images in **A**. **(C-D)** Quantitative RT-PCR analysis of key enzymes of *Ppara* **(C)** and *Ppard* **(D)** in control and *Isg15* KO BMDMs stimulated with or without DMXAA for indicated timepoints (n=4 per group). Data were normalized to 18*S* rRNA. Data represented as means ± SEM. Two-way ANOVA followed by Tukey’s multiple comparisons test. **P* < 0.05; ***P* < 0.01; ****P* < 0.001. n.s., not significant.

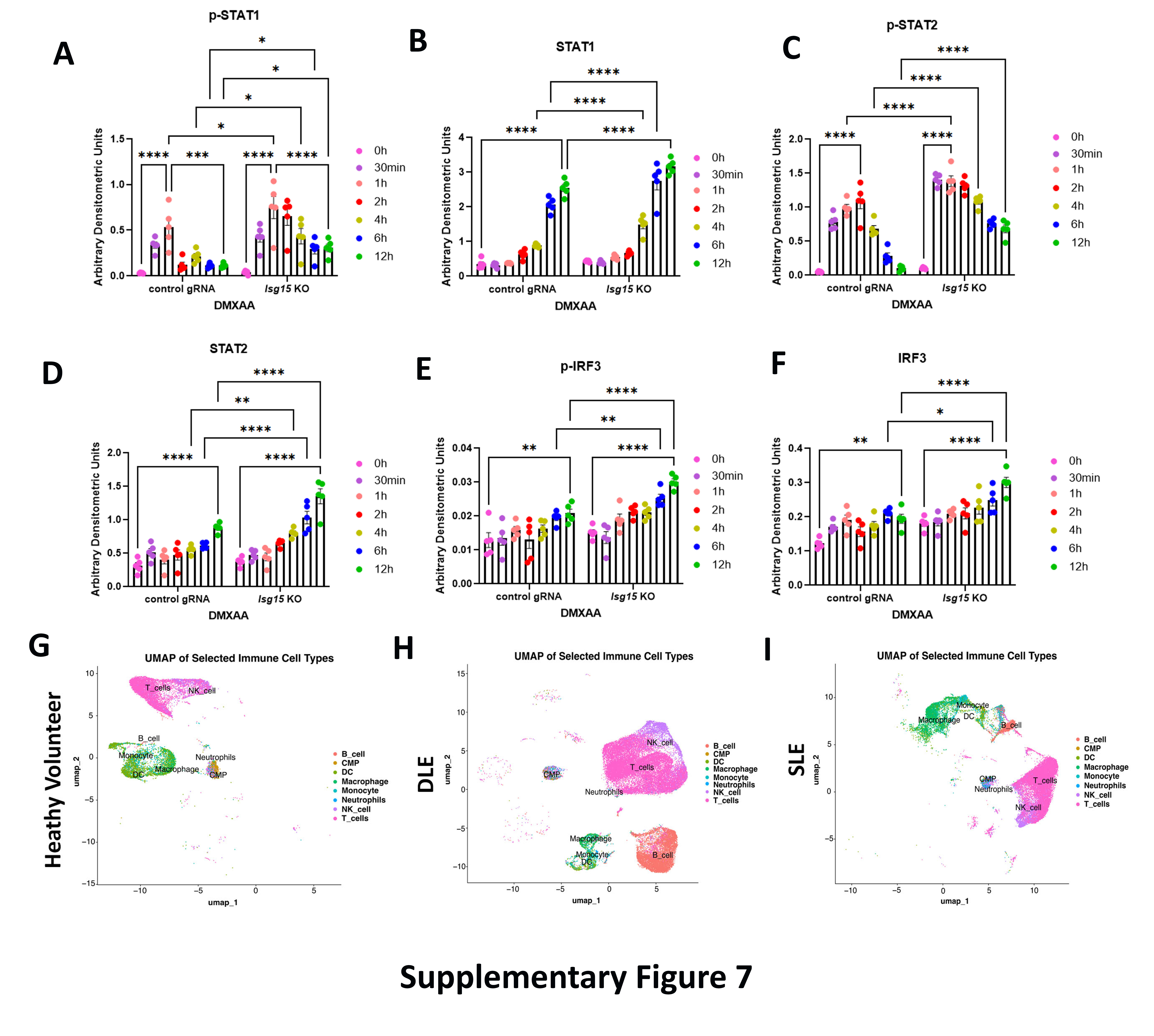

**Supplementary Figure 7: ISG15 negatively regulates T1IFN response**

**(A-F)** Densitometric analysis of immunoblots of p-STAT1 **(A)**, STAT1 **(B)**, p-STAT2 **(C)**, STAT2 **(D)**, p-IRF3 **(E)**, and IRF3 **(F)** relative to vinculin (n=5 per group) in control and *Isg15* KO BMDMs treated with DMXAA at indicated timepoints. **(G-I)** UMAP plot showing the distribution of different immune cell types, including B cells, CMP, dendritic cells (DC), macrophages, monocytes, neutrophils, NK cells, and T cells in dermal tissues of healthy volunteer **(G)**, DLE **(H)** and SLE **(I)** patients.

|  | **Gene** | **Forward Primer** | **Reverse Primer** |
| --- | --- | --- | --- |
| **Mouse** | ***Cpt1a*** | CTCCGCCTGAGCCATGAAG | CACCAGTGATGATGCCATTCT |
| **Mouse** | ***Acat1*** | CAGGAAGTAAGATGCCTGGAAC | TTCACCCCCTTGGATGACATT |
| **Mouse** | ***Hadha*** | TGCATTTGCCGCAGCTTTAC | GTTGGCCCAGATTTCGTTCA |
| **Mouse** | ***Hadhb*** | ACTACATCAAAATGGGCTCTCAG | AGCAGAAATGGAATGCGGACC |
| **Mouse** | ***Acadvl*** | CTACTGTGCTTCAGGGACAAC | CAAAGGACTTCGATTCTGCCC |
| **Mouse** | ***Acadm*** | AGGGTTTAGTTTTGAGTTGACGG | CCCCGCTTTTGTCATATTCCG |
| **Human** | ***CPT1A*** | TCCAGTTGGCTTATCGTGGTG | TCCAGAGTCCGATTGATTTTTGC |
| **Human** | ***HADHA*** | CTGCCCAAAATGGTGGGTGT | GGAGGTTTTAGTCCTGGTCCC |
| **Human** | ***HADHB*** | TACGGGTTTGTTGCATCGGAC | GCCACATTGCTTGTTTTCACTT |
| **Human** | ***ISG15*** | CGCAGATCACCCAGAAGATCG | TTCGTCGCATTTGTCCACCA |
| **Human** | ***ACAT1*** | ATGCCAGTACACTGAATGATGG | GATGCAGCATATACAGGAGCAA |

**Supplementary Table 1: Custom designed primer sets used for quantitative RT-PCR analysis**

|  | **Gene** | **Source** | **Catalogue** |
| --- | --- | --- | --- |
| **Mouse** | ***Ifnb*** | Qiagen | QT00249662 |
| **Mouse** | ***Stat1*** | Qiagen | QT00162183 |
| **Mouse** | ***Stat2*** | Qiagen | QT00160216 |
| **Mouse** | ***Isg15*** | Qiagen | QT00322749 |
| **Mouse** | **18S rRNA** | Qiagen | QT02448075 |
| **Human** | ***IFNB*** | Qiagen | QT00203763 |
| **Human** | **18S rRNA** | Qiagen | QT00199367 |

**Supplementary Table 2: Pre-made and commercially available primer sets used for quantitative RT-PCR analysis**
